# Feedforward mechanisms of masking in mouse V1

**DOI:** 10.1101/2021.04.23.441197

**Authors:** Dylan Barbera, Nicholas J. Priebe, Lindsey L. Glickfeld

**Affiliations:** Center for Learning and Memory, The University of Texas at Austin, 2415 Speedway, Austin, TX 78712, USA; Department of Neurobiology, Duke University Medical Center, Durham, NC 27710, USA

**Author notes:** These authors contributed equally (Nicholas J. Priebe, Lindsey L. Glickfeld). Proofs and Correspondence to and.

**Keywords:** cross-orientation suppression, normalization, plaid, spatial phase, primary visual cortex, LGN, electrophysiology, calcium imaging, optogenetics, parvalbumin, inhibition

## Abstract

Sensory neurons not only encode stimuli that align with their receptive fields but are also modulated by context. For example, the responses of neurons in mouse primary visual cortex (V1) to gratings of their preferred orientation are modulated by the presence of superimposed orthogonal gratings (“plaids”). The effects of this modulation can be diverse: some neurons exhibit cross-orientation suppression while other neurons have larger responses to a plaid than its components. We investigated whether these diverse forms of masking could be explained by a unified circuit mechanism. We report that the suppression of cortical activity does not alter the effects of masking, ruling out cortical mechanisms. Instead, we demonstrate that the heterogeneity of plaid responses is explained by an interaction between stimulus geometry and orientation tuning. Highly selective neurons uniformly exhibit cross-orientation suppression, whereas in weakly-selective neurons masking depends on the spatial configuration of the stimulus, with effects transitioning systematically between suppression and facilitation. Thus, the diverse responses of mouse V1 neurons emerge as a consequence of the spatial structure of the afferent input to V1, with no need to invoke cortical interactions.

## Introduction

As information propagates along the visual pathway, neural circuitry makes fundamental transformations in the representation of the visual scene. There are conflicting views on the roles that feedforward and recurrent cortical circuitry play in generating cortical representations of the visual scene (Carandini & Heeger, 2012; Priebe & Ferster, 2012; Alitto & Dan, 2010). It is clear that the intricate recurrent cortical circuitry modifies representations (Hubel & Wiesel, 1962; Nelson & Frost, 1978; Gilbert & Wiesel, 1990; DeAngelis et al., 1994) but it also clear that the convergent inputs from the thalamus play an essential role in the emergence of cortical selectivity (Chapman et al., 1991; Reid & Alonso, 1995; Ferster et al., 1996; Jin et al., 2011; Lien & Scanziani, 2013). Parsing the contributions of these circuit elements is particularly interesting when multiple stimuli are presented as cortical responses to these stimuli are often nonlinear. One such phenomena is cross-orientation suppression, a form of masking whereby a neuron’s response to an optimally oriented grating is reduced when overlaid by a second grating at an angle other than the preferred orientation, forming a plaid (Blakemore and Tobin 1972; Bonds 1989; Carandini and Heeger 1994; Carandini et al. 1997; DeAngelis et al. 1992; Heeger 1992; Morrone et al. 1982, 1987; Sengpiel et al. 1998).

The orientation specificity of masking has led to mechanistic accounts that rely on interaction between neurons in visual cortex, where orientation selectivity emerges. In cortical accounts, activated pools of cortical neurons mutually suppress one another through recurrent inhibition (Heeger, 1992; Rubin et al. 2015). The presence of two differently oriented stimuli elicits responses (at least transiently) from a larger pool of neurons than either stimulus does alone, resulting in a reduced response. An alternative account considers how masking alters the afferent input to the cortex. The geometry of the plaid stimulus alters the timing and amplitude of thalamic responses: some afferent responses are reduced while others are enhanced. These responses are then transformed by both thalamic and cortical nonlinearities, resulting in suppression (Priebe & Ferster 2006, Li et al., 2006; Koelling et al., 2008).

While cross-orientation suppression is pervasive in cat V1 (DeAngelis et al., 1992; Priebe & Ferster, 2006), the impact of orthogonal stimuli is more heterogeneous in mice and primates (Juavinett & Callaway, 2015; Muir et al., 2015; Muir et al., 2017, Palagina et al., 2017; Ringach et al., 2020; Guan et al., 2020). Although many mouse and primate V1 neurons exhibit robust cross-orientation suppression, other neurons exhibit increased responses when the mask is overlaid. Current accounts for cross-orientation interactions provide explanations for suppression, but not facilitation (Heeger, 1992; Rubin et al., 2015; Priebe & Ferster, 2006; Li et al., 2006; Koelling et al., 2008). The mouse visual system therefore provides an opportunity to address whether a common circuit can explain both cross-orientation suppression and facilitation.

We aim to uncover the mechanisms governing the responses to masking in mouse V1 using *in-vivo* whole cell recordings and two-photon calcium imaging. In line with previous reports, we found a broad range of interactions among cortical pyramidal cells, ranging from cross-orientation suppression to facilitation. We demonstrate that both functional (by sensory adaptation) and optogenetic lesions of V1 activity have little effect on the response profile of V1 neurons to plaids, indicating that a cortical mechanism of masking is unlikely. We also directly assayed how masking manifests in PV+ cortical interneurons and found that they do not contain signals necessary to support both enhancement and suppression. Instead, we demonstrate from a feedforward model that the heterogeneous nature of V1 responses can be explained by the diversity of orientation selectivity in mouse V1. In neurons with high orientation selectivity, effects are dominated by cross-orientation suppression. However, when orientation selectivity is low, responses can be facilitated or suppressed depending on the spatial phase of the plaid. We then demonstrate that by shifting the phase of the plaid, neuronal responses in low orientation-selective neurons systematically shift between cross-orientation facilitation and suppression. Therefore, the diverse responses of mouse V1 neurons to plaid stimuli are succinctly accounted for by spatial structure of the feedforward afferents.

## Results

To measure the interactions between superimposed oriented gratings in mouse V1, we measured GCAMP6 calcium signals from genetically identified pyramidal cells using a two-photon microscope while presenting single drifting gratings of the preferred direction and superimposed orthogonal direction (plaids, **Figure 1A**). As previously reported, neurons varied widely in their responses to plaid stimuli (Juavinett & Callaway, 2015; Muir et al., 2015; Muir et al., 2017, Palagina et al., 2017; Ringach et al., 2020). At one end of the spectrum were neurons whose responses resembled those seen in cats, exhibiting robust decreases in response amplitude to the plaid stimulus compared to their preferred grating alone (**Figure 1B**, left panel). Within the same field of view, other neurons had larger responses to the plaid stimulus than to the single gratings (**Figure 1B**, right panel) or exhibited little, if any, change in response amplitude with the addition of a mask (**Supplementary Figure 1A**).

**Figure 1.**
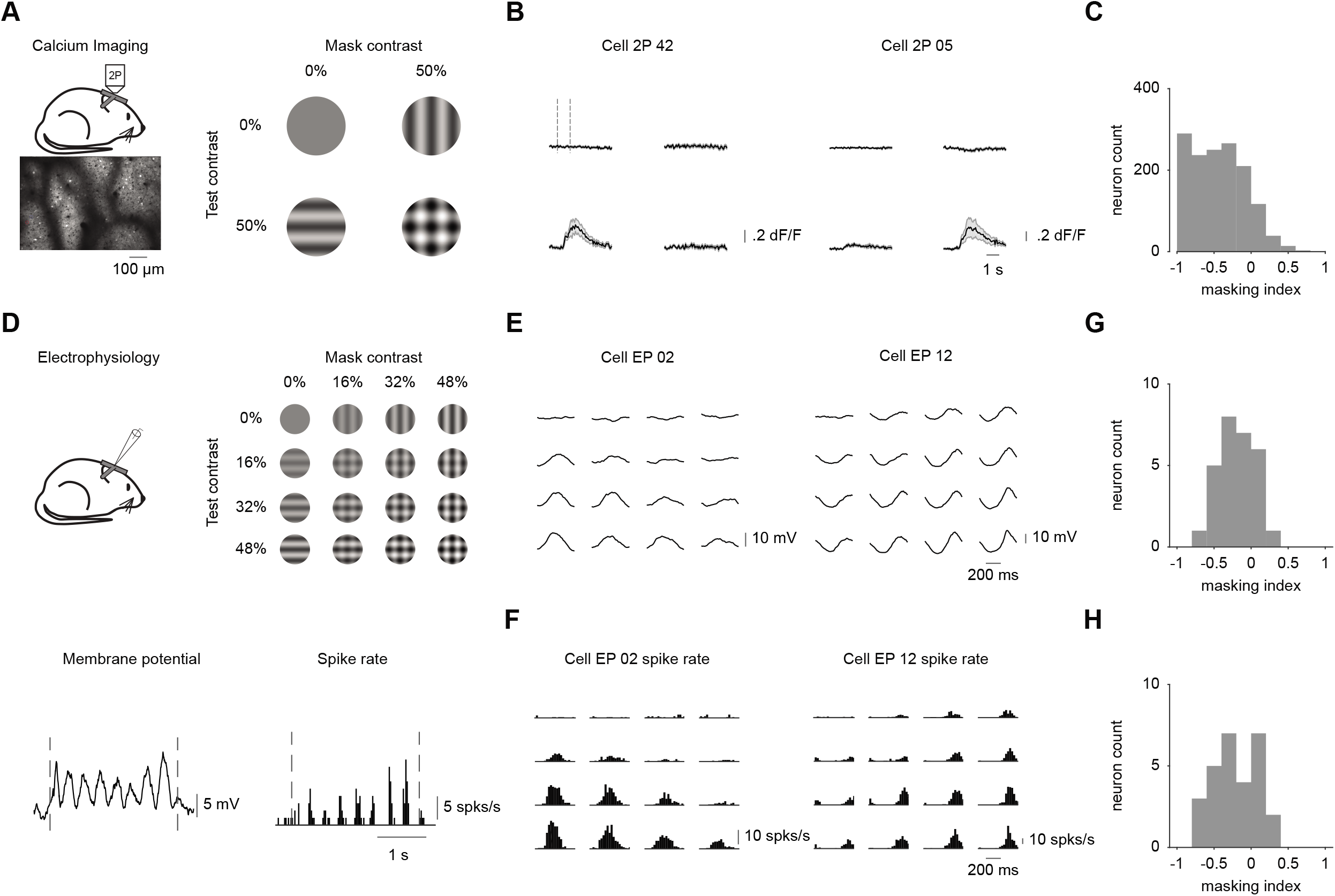
Responses to plaids are diverse in the mouse primary visual cortex. **A**. Left: schematic of imaging setup (top) and example field of view (bottom). Right: stimulus set. **B**. Trial average dF/F responses to gratings and plaids for two example neurons. Shaded error bars are standard error of the mean across trials. **C**. Distribution of masking index for all cells at their preferred direction. **D**. Top: schematic of electrophysiology setup (left) and stimulus set (right). Bottom: trial average responses to a drifting grating for an example neuron for membrane potential (left) and spike rate (right). **E**. Cycle average membrane potential responses to gratings and plaids for two example neurons. **F**. Same as **(E)** for spike rate. **G-H**. Same as **(C)** for membrane potential **(G)** and spike rate **(H)**.

To quantify the interaction between grating components, we computed the masking index (MI, Guan et al., 2020):

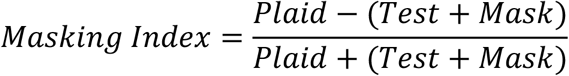

An MI below 0 indicates that the response to the plaid was weaker than that expected by the summed responses to the two grating components (suppression), whereas an MI above 0 indicates facilitation. Although facilitation was present and somewhat common, neurons were on average suppressed in response to plaids (dF/F MI: -0.44 ± 0.01; n = 7 mice, 1426 cells; **Figure 1C**).

We also found diversity in the effects of masking using whole cell recordings in layers II/III and IV (**Figure 1D**). Electrophysiological records of both membrane potential and spiking showed similar patterns of modulations in response to plaids, with examples of both suppression (**Figure 1E-F**, left panel) and facilitation (**Figure 1E-F**, right panel). To quantify these effects, we computed the masking index for the modulation component (F1) of the cycle average (MI computed for highest contrast condition - membrane potential: -0.20 ± 0.04; spike rate: -0.21 ± 0.05; n = 28 cells; **Figure 1G-H**). Of the 28 total neurons in the dataset, 16 were recorded in layers II/III and 12 were from layer IV. There were no significant differences in masking index between the two layers for either membrane potential (unpaired t-test, p=0.50) or spike rate (p=0.26), suggesting that any difference in masking responses between these layers is minor. Masking was evident to a similar degree in the modulation amplitude (F1, see above), mean amplitude (DC, mean membrane potential MI: -0.24 ± 0.06; mean spike rate MI: -0.28 ± 0.05) and their sum (F1+DC, membrane potential MI: -0.21 ± 0.04; spike rate MI: -0.25 ± 0.05) in electrophysiological recordings (**Supplementary Figure 1B-C)**. This similarity is likely the reason that the calcium data, which reflects a transformation of the summed activity, matches well with the electrophysiology data.

Some previous studies have found that facilitation is more common than we report here (Juavinett & Callaway, 2015; Muir et al., 2015; Palagina et al., 2017; Ringach et al., 2019). One possibility is that this may be due to how the orientation of the test and mask gratings are defined. In our dataset, the test stimulus is defined at the preferred orientation. Other studies, however, have not used this criterion and the test stimulus may or may not be at the preferred orientation of the neuron, and this alignment can influence the magnitude of masking (DeAngelis et al., 1992). To determine whether the test orientation impacts the degree of suppression, we computed MI under conditions where the test stimulus was 45 degrees from the preferred orientation (the mask was also rotated by 45 degrees and remained orthogonal to the test stimulus). Under these conditions responses to plaids exhibited significantly less suppression than when tested with the preferred stimulus (MI-dF/F: -0.15 +/- 0.02; n = 526; Wilcoxon rank sum test: p = 4.1e-69; **Supplementary Figure 2**).

One possible explanation for this relationship between masking and stimulus preference could be that neurons’ responses to the preferred stimulus are close to their peak firing rates and therefore cannot exhibit facilitation due to a limited dynamic range. To investigate this, we compared the masking index evoked by plaids composed of gratings at lower contrast. While decreasing the contrast significantly reduced the visually driven response (32% vs 48% contrast-membrane potential: p = 2.6e-4; spike rate: p = 2.1e-4; calcium imaging: p = 5.0e-4), the mean MI did not depend on the contrast of the component gratings (membrane potential: p = 0.58; spike rate: p = 0.73; calcium imaging: p = 0.88). This argues that a limited dynamic range does not explain our data.

### Cortex is not required for masking effects

Two differing mechanistic models have been proposed to account for the effects of masking: the cortical and feedforward models. The cortical model posits that cross-orientation suppression emerges from interactions between distinct populations of orientation tuned cortical neurons (**Figure 2A**). We sought to examine the role of recurrent cortical circuits for both cross-orientation suppression and facilitation by manipulating cortical activity.

**Figure 2.**
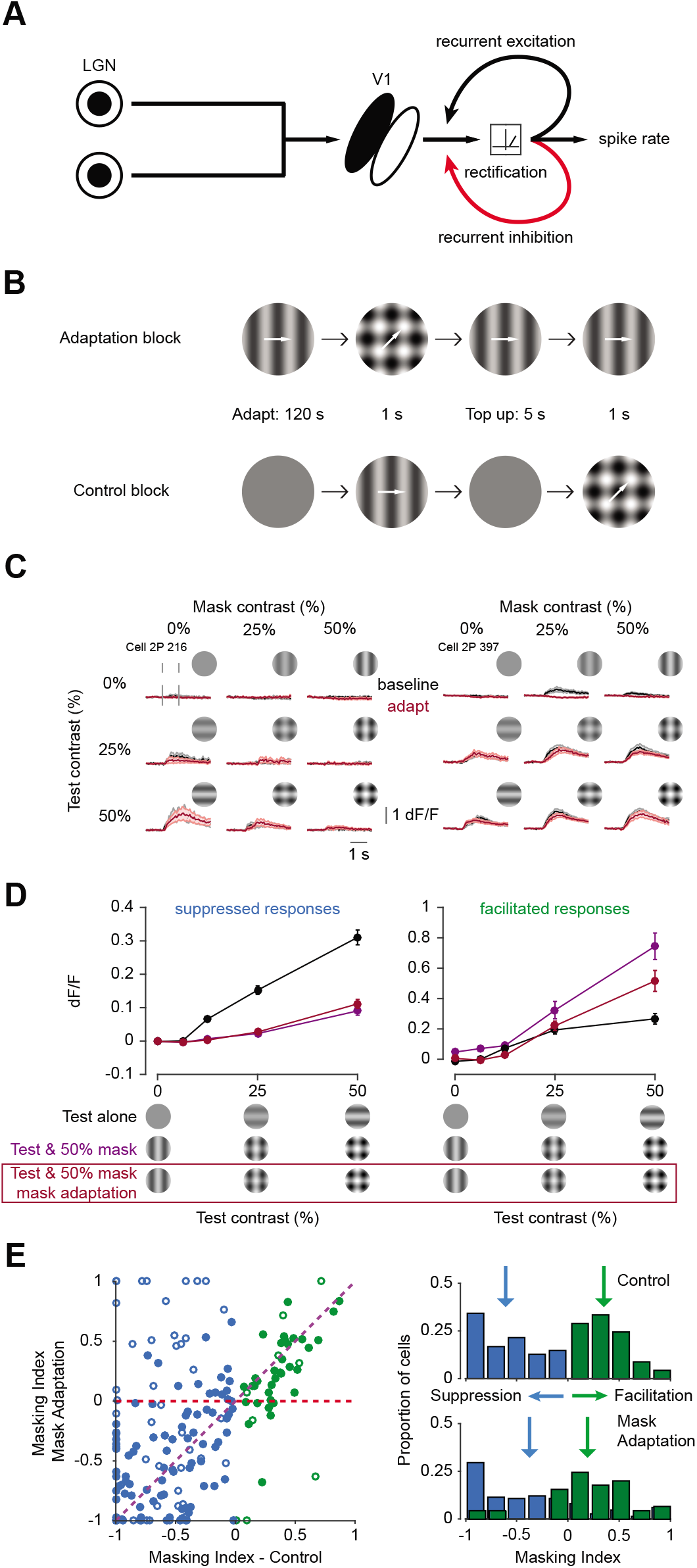
Effects of masking are robust to sensory adaptation **A**. Diagram depicting a circuit for a cortical mechanism of cross-orientation suppression. **B**. Schematic of stimuli for adaptation paradigm (top) and control (bottom). **C**. Example dF/F for two neurons before (black) and after (red) adaptation. Shaded error bars are standard error of the mean across trials. **D**. Average dF/F for test-preferring neurons in response to the test alone (black), test and 50% contrast mask (purple), and test and 50% contrast mask after adaptation (red) for neurons that were suppressed (MI<0; left) or facilitated (MI>0; right) by the mask before adaptation. Error bars are the standard error of the mean across cells. **E**. Left: scatter plot comparing the masking index of all test-preferring neurons before (x-axis) and after (y-axis) adaptation. Color identifies neurons that were suppressed (green) or facilitated (blue) by the mask before adaptation. Filled symbols identify neurons that were significantly suppressed or facilitated before adaptation (unpaired t-test: p<0.05). Red dashed line indicates prediction of cortical model where adaptation should make the masking index zero. Purple dashed line indicates prediction of a non-cortical model where there is no difference between control and adaptation conditions. Right: histograms comparing the masking index before (top) and after (bottom) adaptation. Arrows indicate mean.

We performed two manipulations to weaken the intracortical circuitry. First, we used long term sensory adaptation to functionally lesion cortical activity (**Figure 2B**). Adaptation by prolonged exposure has the effect of reducing responses in V1 while leaving LGN responses largely intact (Sanchez-Vives et al., 2000, King et al., 2016). Therefore, if the effects of masking are invariant to adaptation of one of the component gratings, a cortical mechanism is unlikely (Freeman et al., 2002).

The effects of adaptation on masking were tested in interleaved blocks of adaptation and control trials. In each adaptation block, the mask stimulus (a vertical grating) was initially presented for 120 seconds and then the mask, test (a horizontal grating) and plaid were randomly interleaved between 5 second “top-up” adaptation epochs (**Figure 2B-C**). Control blocks had the same temporal structure, but without stimulus presentation during the adaptation periods. To quantify the effects of adaptation to the mask stimulus on responses to gratings alone, neurons were grouped into two categories based on their responses to gratings: mask preferring and test preferring. In neurons that preferred the mask, sensory adaptation strongly reduced responses to the mask (control dF/F: 0.37 ± 0.03; adapt dF/F: 0.01 ± 0.004; n = 171 cells, 5 mice; two-way ANOVA, main effect of adaptation: p = 3.3e-76; **Supplementary Figure 3A**). This effect did not, however, generalize to responses to the test in test-preferring neurons. Responses in these neurons were only slightly, but still significantly, reduced (control dF/F: 0.3 ± 0.02; adapt dF/F: 0.25 ± 0.02; n = 194 cells, 5 mice; two-way ANOVA, main effect of adaptation: p = 0.002; **Supplementary Figure 3B**). Thus, this manipulation has the effect of selectively reducing cortical responses to one of the two components of the plaid stimulus.

Given that cortical models of suppression depend on the degree to which the cortex responds, selectively weakening responses to the mask stimulus should reduce cross-orientation interactions. However, after adaptation, test-preferring neurons that were suppressed under control conditions generally remained suppressed, and those that were facilitated remained facilitated (Suppressed cells: highest contrast grating dF/F: 0.31 ± 0.02; control plaid: 0.09 ± 0.01; adapt plaid: 0.11 ± 0.01; n = 149 cells; two-way ANOVA, main effect of adaptation: p = 0.27; Facilitated cells: grating dF/F: 0.27 ± 0.03; control plaid: 0.74 ± 0.09; adapt plaid: 0.52 ±0.07; n = 45 cells; two-way ANOVA, main effect of adaptation: p = 0.0001; **Figure 2C-D**).

If cortical circuitry were responsible for the interactions between test and mask stimuli, we expect that the masking index would approach 0 following adaptation (**Figure 2E**, red line). In contrast, if non-cortical mechanisms are responsible for cross-orientation interactions, the masking index should not depend on adaptation (**Figure 2E**, purple line). We found that there was a strong correlation between the masking index before and after adaptation (r = 0.51, n = 189 cells), especially among those cells that remained significantly responsive to the test grating after adaptation (r = 0.76, n = 130 cells). Across the sample population, adaptation resulted in significant changes in the degree of masking (Suppressed cells-control MI: -0.61 ± 0.03; adapt MI: -0.37 ± 0.04; n = 145 cells; paired t-test: p = 3.3e-6; Facilitated cells-control MI: 0.35 ± 0.03; adapt MI: 0.19 ± 0.06; n = 44 cells; p = 0.003), though these effects were less pronounced for the cells that were responsive to the test after adaptation (Suppressed cells: n = 95, p=0.02; Facilitated cells: n = 35, p = 0.006). Despite this regression of the masking index towards zero, both facilitated and suppressed populations nonetheless had masking indexes significantly different than zero after adaptation (Student’s t-test: facilitated-p = 0.003; suppressed-p = 6.2e-13). Thus, cross-orientation interactions remain present after adaptation.

To directly probe the relative contribution of thalamic and cortical circuits to masking we performed whole-cell recordings in layer IV neurons while optogenetically silencing V1. Optogenetic stimulation of inhibitory parvalbumin positive (PV+) interneurons results in robust suppression of cortical pyramidal neurons (Choi & Priebe, 2020; Gu & Cang, 2016; Lien & Scanziani, 2013; Li et al., 2013; Malina et al., 2016). We held neurons at -70 mV using voltage-clamp during optogenetic stimulation and measured the inward currents reflecting the isolated feedforward thalamic input (**Figure 3A**, n=16 neurons).

**Figure 3.**
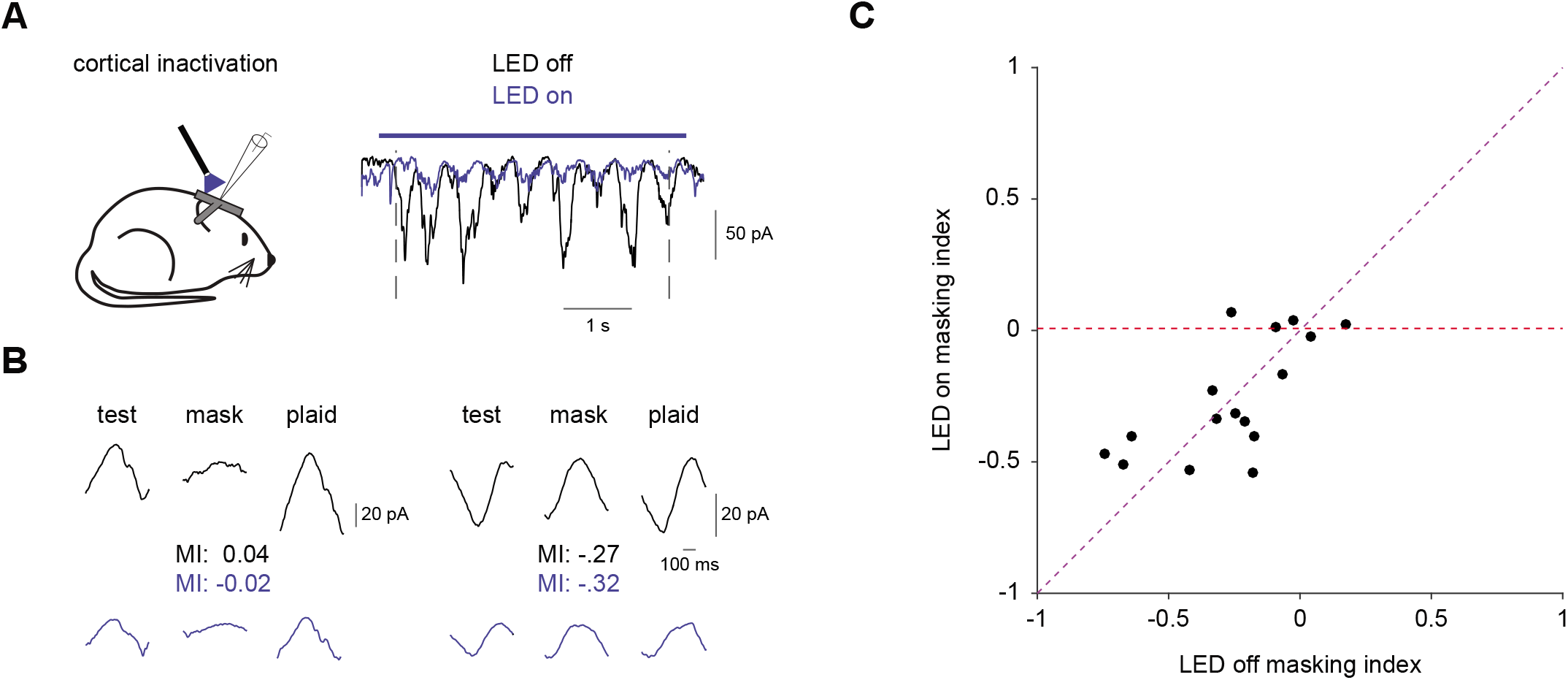
The effects of masking are reflected in the feedforward inputs to V1 **A**. Left: schematic of recording configuration for optogenetic cortical suppression during in-vivo whole-cell recordings. Right: example response to a grating in control (black) and during silencing (blue). **B**. Cycle averages of responses from two example neurons for the test, mask and plaid in control (top) and during silencing (bottom). **C**. Scatter plot comparing the masking index for all cells in control and during silencing. Red dashed line indicates prediction of cortical model where silencing should make the masking index zero. Purple dashed line indicates prediction of a non-cortical model where there is no difference between control and silencing conditions.

Suppression of visual cortex reduces visually-evoked currents (**Figure 3A**). Nonetheless, the amplitude of the isolated thalamic currents were still modulated by the presence of a mask (**Figure 3B**). The mask modulated the response in the same direction (i.e. suppression vs. facilitation) for conditions in which the cortex was active or silenced in all but one neuron (11/12 neurons where -0.1 >MI > 0.1 not included; **Figure 3C**). The magnitude of the response modulation was highly correlated between isolated thalamic and intact cortical circuits (r = 0.67) and there was no significant difference between the average modulation indexes (MI_tot_: -0.22 ± 0.06, MI_thal_: -0.23 ± 0.05; paired t-test: p = 0.88). There were similarly no significant differences between the mean modulation index of the membrane potential records (**Figure 1G**) and both the isolated (unpaired t-test: p = 0.82) and intact (p = 0.65) voltage clamp recordings. The presence of cross-orientation interactions in the isolated thalamic inputs, and their similarity when the cortex is intact, indicates that the seed for cross-orientation interactions lay not in the cortex, but in subcortical structures.

### A feedforward model predicts consequences of masking

Given the limited role of cortical circuits suggested by our experiments, we sought to examine the mechanisms of masking using a feedforward model. In the feedforward model, suppressive interactions stem from nonlinearities (i.e. contrast saturation and rectification) in the thalamic input to the visual cortex (**Figure 4A**). This model has proven successful in explaining cross-orientation suppression in cat primary visual cortex (Priebe and Ferster, 2006; Li et al., 2006). It is not clear, however, whether a feedforward model can reproduce the facilitated responses seen in the mouse.

**Figure 4.**
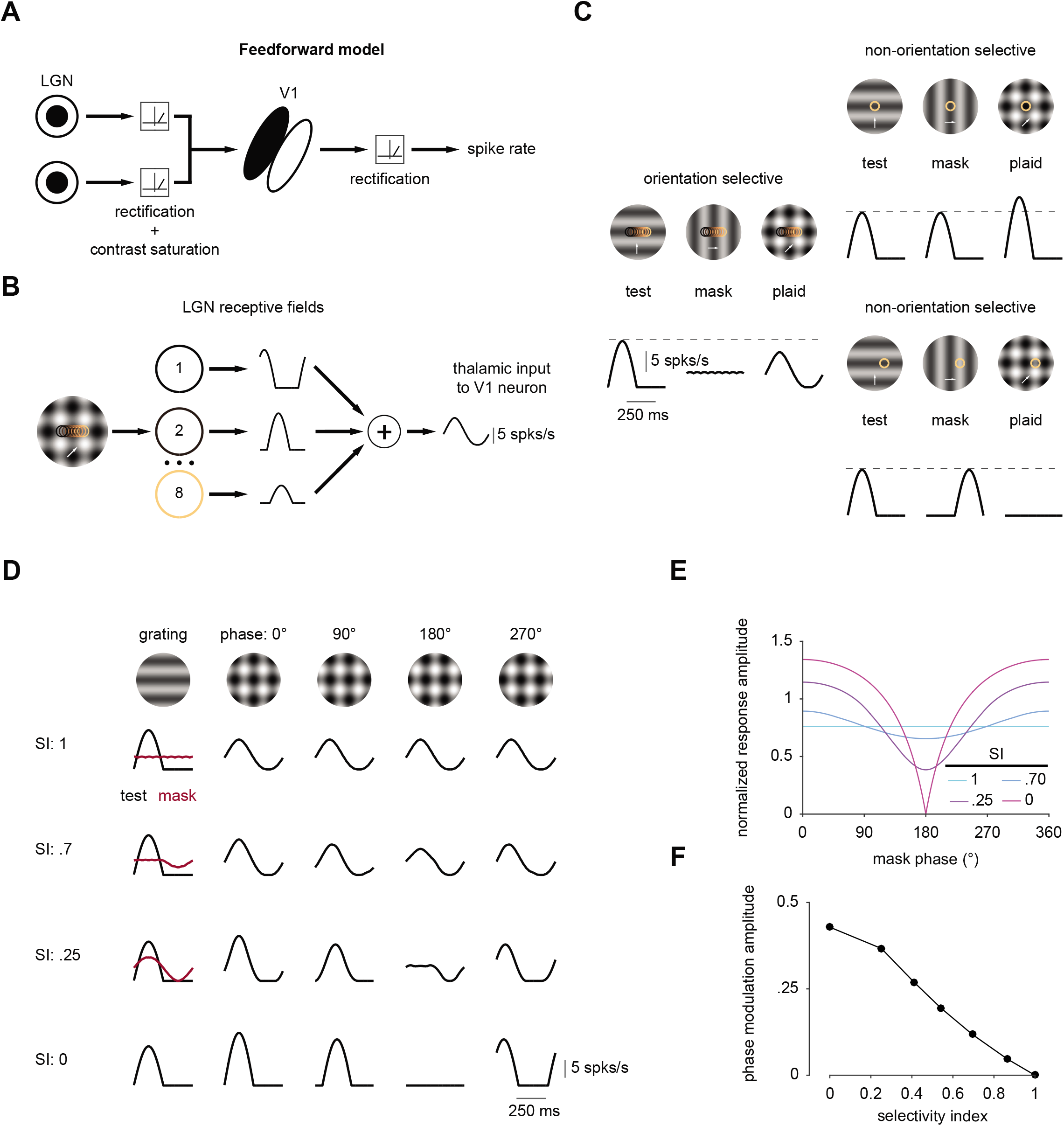
A feedforward model accounts for responses to plaids **A**. Diagram depicting a circuit for a feedforward mechanism of cross-orientation suppression. **B**. Framework for the implementation of the feedforward model. Each circle represents the receptive field of an individual thalamic afferent. Traces reflect responses of the afferents to one stimulus cycle. **C**. Model responses to the test, mask and plaid for an orientation selective (left) and two non-selective (right) model neurons with different receptive field positions (yellow circle). **D**. Model neuron responses to test (black, first column), mask (red, first column) and plaids at multiple stimulus phases (columns 2-5). Each row corresponds to a different selectivity for the test and mask as measured by the selectivity index (SI). **E**. Response amplitude to the plaid stimulus as a function of stimulus phases for cells with varying selectivity index. **F**. Phase modulation amplitude as a function of selectivity index for model neurons.

In the feedforward model, input from thalamic responses to grating and plaid stimuli are generated based on measured contrast and rectification nonlinearities (Priebe & Ferster, 2006). Thalamic relay cells with spatially offset receptive fields converge onto cortical neurons as a basis for their orientation selectivity (Hubel & Wiesel, 1962). This precise organization leads to thalamic responses that are temporally aligned when a preferred orientation grating is presented, resulting in a large modulation of the aggregate input, which is analogous to membrane potential (**Figure 4B**).

Mouse V1 neurons, however, exhibit a broad range of orientation selectivities (Tan et al., 2011; Scholl et al., 2013). Many neurons are only weakly orientation tuned, lacking a precise alignment of their thalamic afferents. In some neurons the response to the orthogonal gratings may be nearly as large as the response to the preferred orientation (Jia et al., 2010). We therefore constructed a feedforward model in which the degree of orientation selectivity was adjusted by controlling the spatial offset of thalamic afferents. When selectivity is altered in this manner, the predicted aggregate thalamic response to a plaid stimulus varied: it could either show suppression or facilitation relative to the response to a single grating (**Figure 4C**).

Notably, we found that this simple model predicts that responses of weakly tuned neurons to plaids are sensitive to the precise location of their receptive field relative to the stimulus (**Figure 4C**, right panel). For example, if the receptive fields of thalamic neurons are positioned at plaid locations where large excursions of luminance occur (**Figure 4C**, top right), model neurons exhibit large responses. If, on the other hand, the receptive fields are positioned at a location for which the mask reduces luminance modulations (**Figure 4C**, bottom right), model neurons exhibit little, if any, response. By comparison, orientation selective model neurons average across all of these regions, and therefore changing the receptive field positions of these cells results only in timing changes.

To explore this interaction between receptive field location and orientation selectivity on the effect of masking, we systematically varied two model features: the degree of stimulus selectivity and the spatial phase of the mask grating (**Figure 4D**). As receptive field locations are fixed, shifting the spatial phase of the mask effectively alters the relative positions of the luminance excursions and receptive field. We quantified the degree of selectivity of model neurons by computing the selectivity index (SI):

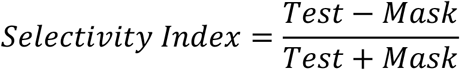

Altering the spatial phase of the mask stimulus had little impact on evoked responses of highly selective neurons: all mask spatial phases resulted in suppression. However, as selectivity index declines, the mask spatial phase alters both the amplitude and timing of plaid responses. These modulations were systematic, with the largest facilitation occurring when the component gratings evoked responses that are spatially, and therefore temporally, in phase (**Figure 4E**). The sensitivity of each cell to the mask spatial phase is summarized by its spatial phase modulation amplitude. Notably, the model predicts a strong inverse relationship between modulation amplitude and stimulus selectivity (**Figure 4F**).

### Spatial phase modulates the effect of masking

To test the feedforward model prediction that responses are sensitive to mask phase, we measured V1 responses to plaids at multiple spatial phases using whole-cell recordings and calcium imaging. We presented plaids composed of a test grating whose phase was fixed while varying the phase of the mask grating at four or eight different phases. Varying the stimulus phase led to dramatic shifts in the neuronal response amplitude to plaids, often shifting between suppression and facilitation in a single neuron (**Figure 5A-C**, left panel).

**Figure 5.**
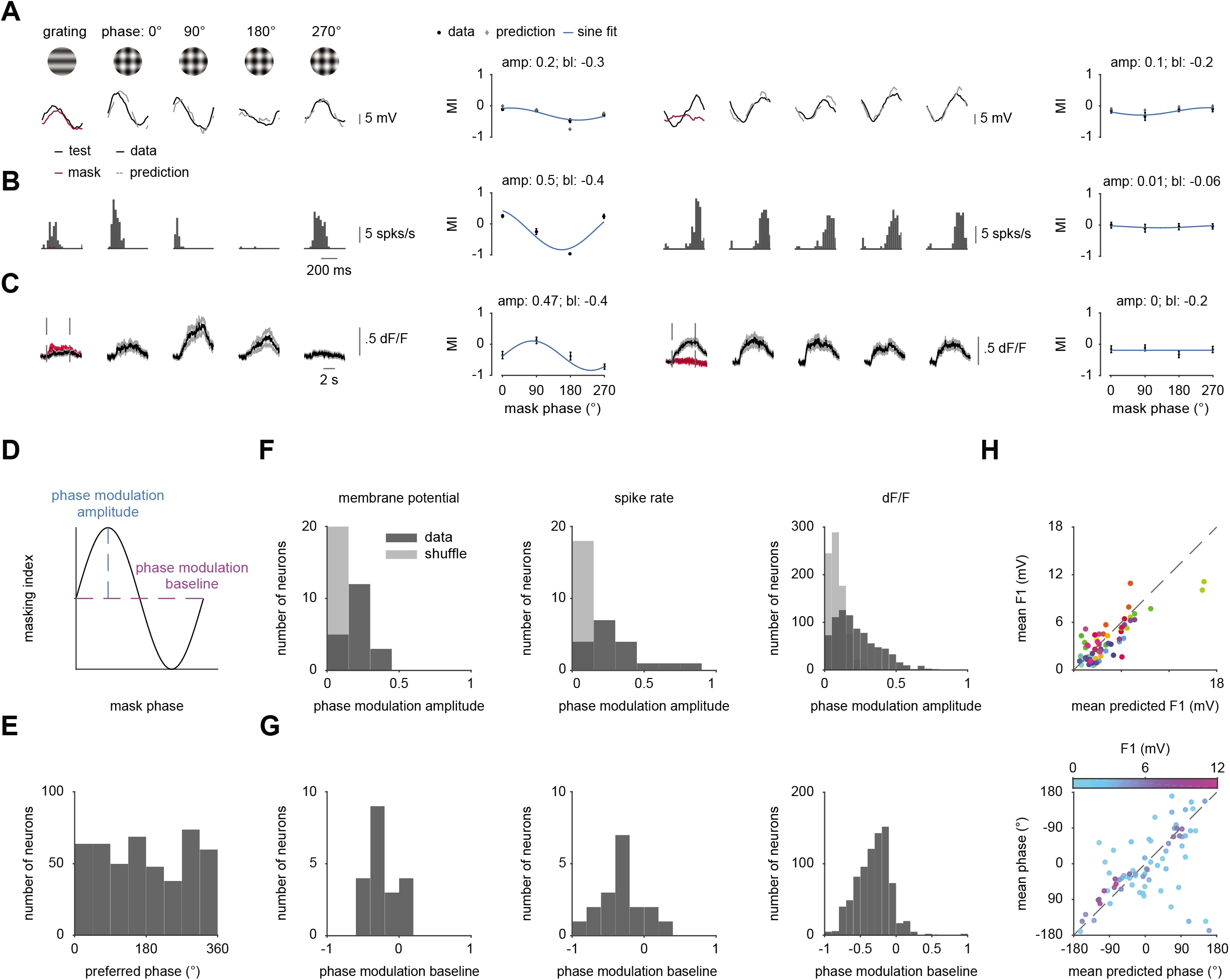
Responses to plaids are phase sensitive **A**. Cycle-average membrane potential responses to test (black), mask (red) and plaids at multiple stimulus phases for two example neurons. Gray dashed lines indicate a linear prediction based on the response to the test and mask alone. Plots depict the masking index measured at four stimulus phases (black) and fit with a sinusoid (blue; amp-amplitude; bl-baseline). Gray dots illustrate linear prediction. **B-C**. Same as **(A)** for spike rate **(B)** and dF/F **(C). D**. Diagram depicting the phase modulation amplitude and baseline of the sine fit. **E**. Histogram of the preferred stimulus phase in the imaging dataset. **F**. Histograms showing the phase modulation amplitudes for membrane potential (left), spike rate (center) and dF/F (right). Light bins depict a shuffled control. **G**. Same as **(F)** but for phase modulation baseline. **H**. Scatter plots comparing the amplitude (top) and phase (bottom) of the linear prediction (x-axis) and membrane potential response (y-axis) to plaids across phases. Four responses are plotted per neuron to represent the response to each stimulus phase. In top panel, color code is by neuron; in bottom panel, color code is by F1 modulation amplitude.

Across our electrophysiological and imaging datasets we found that the spatial phase of the mask stimulus elicits large, significant changes that follow the expectations of the feedforward model. In our electrophysiological records 65% (13/20 cells, by membrane potential) of neurons were significantly modulated by stimulus phase (one-way ANOVA: p < 0.05). A similar proportion of neurons were significantly modulated by phase in our calcium imaging dataset (57%, 468/814 cells, 11 mice). To quantify these phase-dependent interactions, we fit the masking index at each stimulus phase with a sinusoidal function and computed both the amplitude and baseline of the sine-wave fit (**Figure 5D**). The amplitude indicates the phase sensitivity of that neuron while the baseline indicates whether the neuron is facilitated or suppressed across phases. Neurons exhibited a high degree of phase sensitivity across the population (**Figure 5F**) and the phase modulation amplitude was significantly greater than shuffled controls for membrane potential (median sine amplitude: 0.21, shuffle: 0.006; Wilcoxon signed rank: p<0.001), spike rate (median sine amplitude: 0.22, shuffle: 0.002; p<0.001) and dF/F (median sine amplitude: 0.19, shuffle: 0.08; p <0.001). Phase modulation baseline was negative on average (membrane potential: -0.26 ± 0.04; spike rate: -0.31 ± 0.07; dF/F: -0.31 ± 0.01 for dF/F), indicating that neurons were more likely to be suppressed by the plaid across phases (**Figure 5G**). Altogether, this reveals a strong dependence of stimulus phase in the effect of masking, consistent with the feedforward but not the cortical model.

What determines when a given stimulus phase will elicit a large or small response? In the model, the receptive fields are fixed at a particular point in the plaid. This generates the maximum response when the two component gratings are aligned in phase. In our recordings, we do not have access to the location of the individual LGN receptive fields. Therefore, the plaid phase that elicits the maximum response is uniformly distributed (**Figure 5E**).

Whereas the preferred mask phase is uniformly distributed, the feedforward model predicts that the phases at which suppression and enhancement occur depend on the timing of the responses to the test and mask alone. Specifically, there should be enhancement when those responses are temporally aligned and suppression when they are out of phase (**Figure 4D-E**). To test whether this linear combination of the test and mask can account for the amplitude and timing of responses to the plaid, we summed the membrane potential responses to the test and mask alone and after shifting the timing of the mask response in accordance with the expected change in phase (**Figure 5A**, dotted lines). This linear computation was highly predictive of both amplitude (r = 0.82) and timing (r = 0.53) of the response to the plaid (**Figure 5H**). Thus, the phase dependence of masking is directly predicted by the linear summation of LGN inputs in V1 neurons.

### Spatial phase modulation depends on stimulus selectivity

In addition to predicting that masking will be sensitive to the stimulus phase, the feedforward model predicts a nearly linear relationship between selectivity index and phase modulation amplitude (**Figure 4F**). Upon initial inspection, however, the membrane potential responses only partially follow this pattern (**Figure 6A**). Neurons with high selectivity indices exhibit membrane potential responses that conform to the predicted relationship, with a nearly linear pattern of decreasing phase modulation amplitude as selectivity increases. Neurons with low selectivity, however, exhibit both low and high phase modulation amplitudes, contrary to the expectations of the feedforward model.

**Figure 6.**
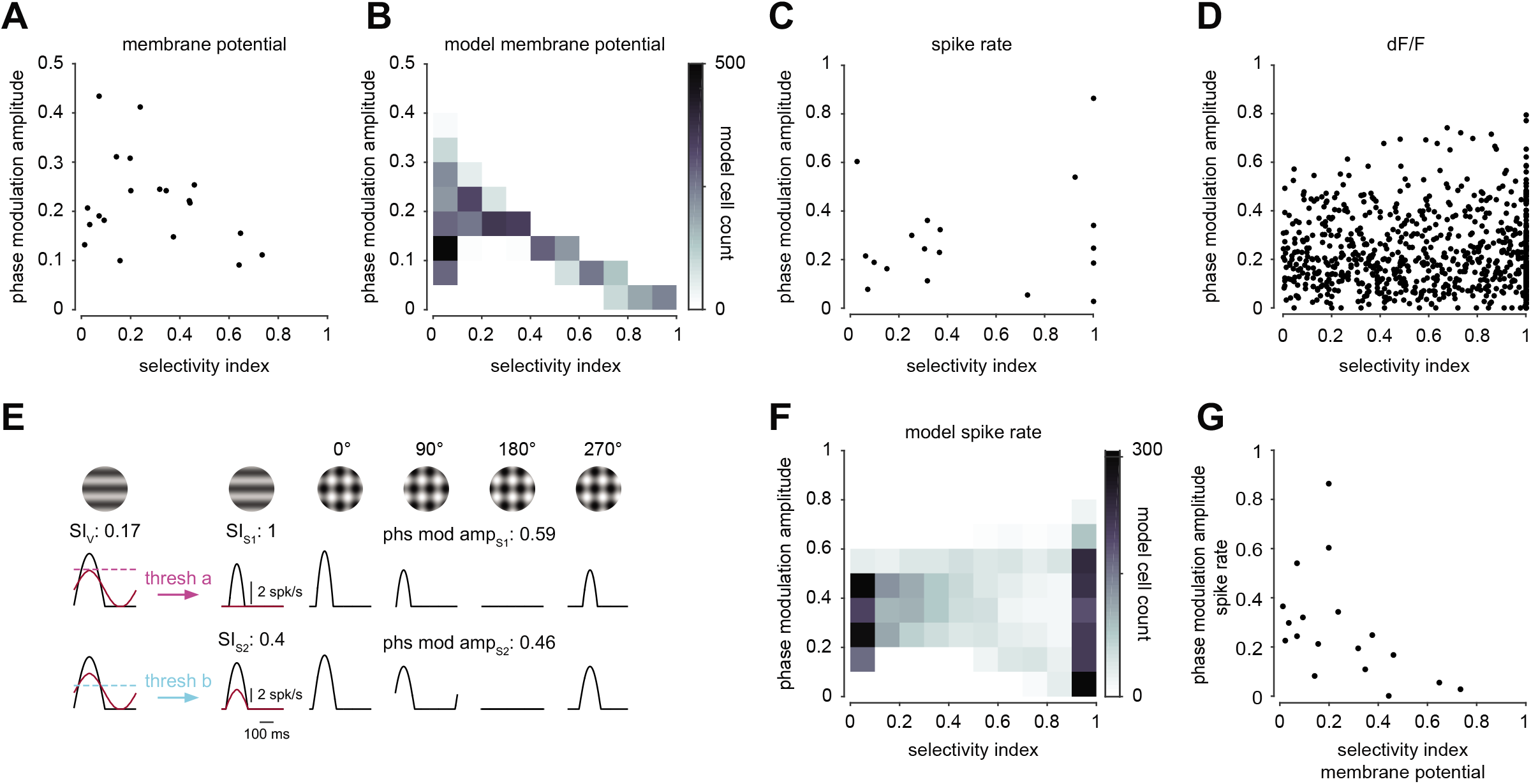
Phase sensitivity depends on the selectivity of membrane potential responses. **A**. Scatter plot comparing the selectivity index (x-axis) and phase modulation amplitude (y-axis) of membrane potential responses. **B**. Heatmap comparing selectivity index and phase modulation amplitude for a simulated population of model neurons sampling at only four phases. **C**. Same as **(A)** for spike rate. **D**. same as **(A, C)** for dF/F. **E**. Responses of a model neuron with two different thresholds demonstrating the effect of threshold on SI and phase modulation amplitude. **F**. Same as **(B)** for spike rate. **G**. Scatter plot comparing membrane potential SI to spike rate phase modulation amplitude.

What can account for this apparent contradiction between the model and data? One difference between the data and model concerns the number of stimulus phases used to measure phase modulation amplitude. Model neuron responses were measured at 360 stimulus phases. Due to experimental constraints, membrane potential data for most neurons were gathered at only 4 stimulus phases. Thus, we hypothesized that the under-sampling of stimulus phase in our dataset might result in an underestimate of phase modulation by missing points that reflect the peak or trough of the function.

To determine whether our experimental under-sampling of the full phase tuning curve might be biasing our measurements, we restricted our model to only sample the limited set of phases explored in the experiment. Under these conditions, the model predicts a linear relationship at higher SI’s and a scattering of phase modulation amplitude values at lower SI’s, matching the experimental data (**Figure 6B**).

While plaid phase sensitivity and selectivity are strongly linked at the membrane potential level, there is no clear relationship at the spiking level across the population of the same neurons (**Figure 6C**), or in our calcium imaging dataset (**Figure 6D**). We hypothesized that this discrepancy lies in the nonlinearities (and the variation of these nonlinearities across neurons) that define how membrane potential is converted to spikes. To test this idea, we passed model responses through a rectifying nonlinearity which varies in threshold (**Figure 6E**). This variation in threshold captures the weak relationship between spike rate SI and phase modulation amplitude (**Figure 6F**). This is due mostly to the effects of threshold on selectivity index which is significantly altered by the threshold nonlinearity (ephys: Vm SI versus spike rate SI paired t-test; p = 0.01) while the phase modulation amplitude is comparatively robust to threshold (no significant difference, p = 0.27). Because the phase modulation amplitude is robust to threshold differences, it is strongly related to the membrane potential selectivity index both in our data (**Figure 6G)** and the feedforward model (**Supplementary Figure 4A-B**). These findings highlight how an intervening nonlinearity, like spike threshold, can obscure a strong relationship between parameters at the membrane potential level.

In contrast to phase modulation amplitude, phase modulation baseline exhibited a slight positive correlation with membrane potential selectivity index (r = 0.3; **Supplementary Figure 4C-D**). This relationship was also altered by threshold, as selectivity index and phase modulation baseline exhibited a small negative correlation in both spike rate (r = -0.24; **Supplementary Figure 4C**) and calcium imaging (r = -0.2; **Supplementary Figure 5A-B**). Taken together, these results suggest that as selectivity increases, the amplitude of the membrane potential phase modulation curve decreases while the mean of the curve shifts upwards, resulting in a neuronal response that is less sensitive to phase and on average less modulated by the mask. These dependencies emerge from a simple feedforward model that includes contrast saturation and response rectification.

### Cross-orientation interactions manifest differently in PV+ interneurons than pyramidal cells

The suppressive and facilitative effects of mask stimulation emerge naturally in a feedforward model, but V1 neurons also receive extensive intracortical inhibitory inputs that may play a role in this process. We therefore sought to examine the impact of plaid stimulation on a prominent subtype of cortical interneuron: PV+ neurons. PV+ interneurons are broadly-selective and their activation linearly suppresses pyramidal cell firing, implicating them in normalization processes such as cross-orientation suppression (Sohya et al. 2007; Niell and Stryker, 2008; Kerlin et al., 2010; Ma et al., 2010; Runyan et al., 2010; Hofer et al., 2011; Atallah et al., 2012; Wilson et al., 2012). PV+ interneurons were selectively labeled using *PV*-Cre knock-in mice injected with flex-GCaMP6s and we used two photon calcium imaging to measure responses to gratings and plaids at multiple spatial phases (**Figure 7A-B**). We then compared these measurements to responses obtained from the population of genetically identified pyramidal neurons described in **Figures 5-6**. Our expectations were that if PV+ cells contribute to the overall suppression of pyramidal cells then plaid stimuli should provide weaker suppression (as predicted by the supralinear stabilized network (SSN) model (Ozeki et al., 2009; Rubin et al., 2015; Ahmadian et al. 2013)) or even facilitation (as predicted by the normalization model (Heeger et al., 1992)). Further, if PV+ cells are responsible for the phase-dependent modulation of pyramidal cells then their suppression should be as modulated as pyramidal cells.

**Figure 7.**
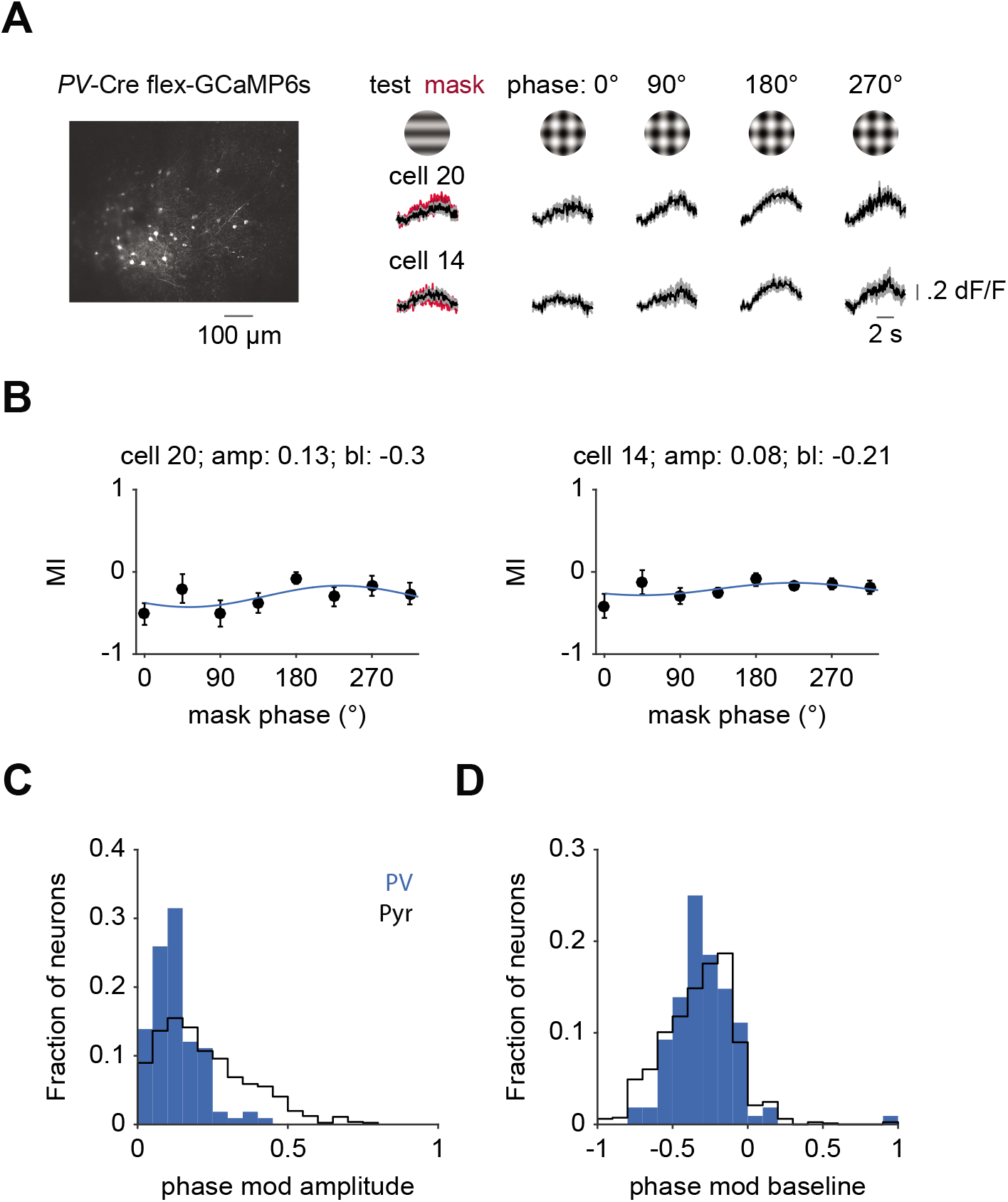
Limited phase sensitivity in inhibitory PV+ neurons **A**. Left: example field of view of a PV-Cre mouse injected with flex-GCaMP6s. Right: example responses of two Pv+ neurons to gratings and plaids at multiple stimulus phases. **B**. Plots depicting the masking index of the two example neurons in (**A**) at eight stimulus phases. Data (black) are fit with sinusoids (blue) to measure the baseline and amplitude of the modulation. **C**. Histogram showing distributions of phase modulation amplitude for PV+ (filled, blue) and excitatory (outline, black) neurons. **D**. Same as (**C**) for phase modulation baseline.

Consistent with the literature, PV+ cells were significantly less selective than pyramidal cells, whether quantified across orientations (mean OSI-PV: 0.39 ± 0.03, n = 108 cells, 5 mice; Pyr: 0.75 ± 0.01, n = 281 cells, 7 mice; p<2.88e-21, Mann-Whitney U-test) or by the selectivity for the test and mask gratings (SI-PV: 0.30 ± 0.02, n = 108 cells, 5 mice; Pyr: 0.61 ± 0.01, n = 814 cells, 11 mice; p = 9.56e-19). We measured the effects of masking at multiple mask spatial phases to extract both how the specific geometry of the visual stimulus modulates inhibitory cell responses in addition to the baseline degree of masking (**Figure 7B**). Whereas mask phase has a profound impact on pyramidal cell responses, significantly fewer PV+ cells were modulated by mask phase (22/108; p = 3.75e-13, Chi-squared test). Because response amplitude varied weakly across mask spatial phase, the masking index exhibited only weak modulations (median amplitude-PV: 0.11; p < 3.81e-12, Mann-Whitney U-test; **Figure 7C**). In contrast, we found that pyramidal cells and PV+ interneurons exhibited similar degrees of overall suppression across mask phases (mean baseline-PV: -0.28 ± 0.02; p = 0.46, Mann-Whitney U-test; **Figure 7D**). The convergence of both of these results indicate that the PV+ interneurons respond to distinct aspects of the visual plaid than their pyramidal cell counterparts. Further, PV+ neurons do not exhibit responses that could account for the spatial phase dependent masking or have weaker suppression than pyramidal cells.

## Discussion

The feedforward model provides a simple mechanistic account of masking in the primary visual cortex. Both cross-orientation suppression and facilitation arise from the same feedforward mechanism, with response diversity accounted for by the broad range of orientation selectivities in mouse V1. We have demonstrated that the spatial phase of the plaid is a determining factor of response amplitude of V1 neurons with low selectivity between the test and mask stimulus. This phase sensitivity naturally emerges from the spatial structure of the afferent receptive field, and does not rely on cortical interactions.

These results point to two factors which determine the strength of neuronal response to plaid stimuli: 1) the selectivity of the V1 neuron for the test over the mask and 2) the relative position of the thalamic afferents and the luminance excursions of plaid stimuli (i.e. spatial phase). When neurons are highly selective for the test over the mask, a superimposed mask results in cross-orientation suppression due to contrast saturation and spike rectification. If selectivity is low, however, the neurons can be shifted systematically between facilitation and suppression by adjusting the relative phase of the test and mask. In light of this sensitivity, which depends on the stimulus, not on the cell, it does not seem appropriate to categorize V1 neurons as “plaid suppressed” or “plaid preferring” on the basis of a facilitated response to a plaid at one spatial configuration (Guan et al, 2020). Furthermore, experiments which measure responses to plaids in mouse V1 must consider the effect of spatial phase. For example, plaids have been used to measure selectivity to complex motion patterns in mouse V1 (Juavinett & Callaway, 2015; Muir et al., 2015; Muir et al., 2017; Palagina et al., 2017) and the reported effects might be reconsidered in light of our results. We have likely underestimated the total range of phase-induced modulations across the population as we employed a limited number of spatial phases which may not match the peaks or troughs of the phase interactions (**Figure 6 A**,**B**). It is likely that there are additional stimulus features not explored here, such as spatial frequency, which contribute to the pattern of responses as well (DeAngelis et al., 1992).

By reversibly lesioning V1 with both sensory adaptation and optogenetics, we have shown that there is no need to invoke a cortical mechanism to explain the effects of masking in mouse V1. This explanation is inconsistent with circuit models such as the SSN model which relies on intracortical inhibition to explain cross-orientation suppression (Rubin et al., 2015). Inhibitory interneurons, however, do not have a receptive field geometry that could render them sensitive to the relative phase of the test and the mask (Niell & Stryker, 2008; Ma et al., 2010; Liu et al., 2009). Our recordings from PV+ interneurons reflect this difference in functional selectivity, as these neurons were significantly less modulated by the spatial phase of the mask than excitatory neurons. Moreover, the SSN model proposes to account for both surround suppression and cross orientation suppression using the same circuit motif. The mechanisms underlying these phenomena, however, are likely separate, as the dynamics of surround suppression and cross-orientation interactions are distinct (Smith, Bair & Movshon 2006), and the effects of surround stimuli do not relieve cross-orientation interactions (Trott and Born 2015).

Our experiments were confined to mouse V1, but there is reason to believe that the findings presented here are applicable to other species. Previous research has demonstrated that in both cats and primates feedforward mechanisms account for the effects of cross-orientation suppression in V1 (Priebe & Ferster, 2006; Li et al., 2006, Koelling et al., 2008, Freeman et al., 2002). Our finding that the masking index (and its phase sensitivity) depends on the relative response to the test and the mask also resolves the reported discrepancies across species in the effects of masking. While the diversity of orientation tuning is largest in mouse V1, primate V1 does contain neurons with low orientation selectivity (Ringach et al., 2002; Tan et al., 2014), whereas neurons in cat V1 are uniformly sharply tuned (Scholl et al. 2013). Thus, this explains the frequency of observations of facilitation being highest in the mouse (Juavinett & Callaway, 2015; Muir et al., 2015; Muir et al., 2017; Palagina et al., 2017), present in the primate (Guan et al., 2020), and absent in the cat (DeAngelis et al. 1992, Priebe & Ferster 2006). Finally, our work resolves that differences in the prevalence of facilitation and suppression across mouse studies lies in the definition of the test stimulus. When the test stimulus is defined as the preferred orientation greater suppression is observed than when the test is defined as nonpreferred-orientations.

In our model, lateral geniculate receptive fields were imagined as circularly symmetric, lacking orientation selectivity. Some mouse LGN neurons, however, have been shown to exhibit a limited degree of orientation selectivity (Piscopo et al, 2013; Scholl et al., 2013; Zhao et al., 2013; Suresh et al., 2016). It is possible that these orientation-selective LGN neurons respond in a way not accounted for by our simple model. Such elaborations suggest that some of the nonlinear interactions we have explored in the feedforward inputs may arise as early as the retina. In addition, we modelled thalamic subfields as spatially aligned along the preferred orientation axis. Orientation selectivity in mouse V1, however, may emerge from alternative thalamic alignments, including randomly distributed feedforward inputs (Pattadkal et al., 2018) or offset ON and OFF thalamic inputs (Liu et al. 2010; Lien & Scanziani, 2013). These distinct receptive field models, however, all have thalamic inputs that exhibit nonlinearities and geometries that will interact with plaids in a similar fashion as our simple model, and thus are unlikely to substantially change our predictions. Finally, our model is based on the integration of LGN inputs in cortical cells, typically thought to occur primarily in layer IV. However, we observed comparable effects of masking among electrophysiological records from layers 4 and II/III (and comparable results from calcium imaging from layer II/III). This may reflect the promiscuous input from the LGN to both cortical layers (Ji et al, 2016; Kondo & Ohki, 2016). Alternatively, this may reflect precise retinotopic connectivity that preserves the geometric interactions between test and mask across multiple synaptic connections (Cossell et al. 2015).

The behavior of PV+ neurons seems to be at odds with our feedforward model. The feedforward model predicts that the degree of phase modulation is inversely related to orientation tuning. Like others, we found that PV+ neurons are weakly tuned for orientation (Niell and Stryker, 2008; Ma et al. 2010; Sohya et al. 2007; Kerlin et al., 2010; Atallah et al., 2012), but they were only weakly modulated by mask spatial phase. This discrepancy may lie in how PV+ neuron responses emerge. Unlike pyramidal cells, PV+ neurons integrate the responses of nearby neurons with distinct spatial receptive fields and orientation preferences, which would blur the impact of mask spatial phase (Kerlin et al., 2010; Hofer et al., 2011; Scholl et al., 2015; Runyan and Sur, 2013; Bock et al. 2011). While PV+ neurons may not provide the input necessary to account for the spatially-specific modulations we describe here, they may nonetheless acts as local stimulus energy sensors that provide a normalization signal for visual cortex. It is important to note, however, that the SSN model proposed that while both excitatory and inhibitory cells exhibit suppressive effects, the effects should be weaker in inhibitory cells than pyramidal cells (Rubin et al. 2015). We have found instead that the overall degree of suppression in PV+ inhibitory cells matches that of pyramidal cells.

In contrast to circuit models, other cortical models offer a computational perspective on cross-orientation suppression. In the normalization model, for example, the two different stimulus orientations composing the plaid lead to the activation of a greater number of cortical neurons, resulting in increased suppression (Heeger et al., 1992). Although that circuit is not consistent with our proposed mechanism, the normalization model provides a compact description of many of the nonlinear response properties of cortical neurons and is a powerful conceptual model of cortical responses (Carandini and Heeger 2011). One interpretation of our results is that the normalization model provides general rules that govern cortical responses and in the case of cross-orientation suppression, the feedforward circuit implements those rules.

One successful approach to understanding sensory processing is to focus on the computation being performed separate from the specific implementation (Marr, 1982; Marr and Poggio 1976). As shown here, however, neurons may respond in ways that are superficially misleading. For example, one might interpret cross-orientation facilitation as a unique computation, separate from that of cross-orientation suppression. By taking the alternate approach of focusing on the mechanistic details we have revealed that cross-orientation interactions are a direct consequence of the spatial geometry and position of the plaid. Nonetheless, both the computational and mechanistic approaches - and the interactions between these approaches - are essential to understand sensory processes.

## Acknowledgments

We thank Grace Link for surgical assistance and both Hillel Adesnik and Uday Jagadisan for their valuable feedback on a previous version of the manuscript. This work was supported by grants from the National Eye Institute (1R01-EY031328 to L.L.G. and R01-EY025102 to N.J.P).

## Author Contributions

Conceptualization: D.B., N.J.P, L.L.G.; Methodology: D.B., N.J.P, L.L.G.; Investigation: D.B., L.L.G. Formal Analysis: D.B., N.J.P, L.L.G.; Writing - Original Draft: D.B.; Writing - Review and Editing: D.B., N.J.P, L.L.G.; Funding Acquisition: N.J.P, L.L.G.

## Methods

### Animals

#### Electrophysiology

Experiments were conducted using adult C57/BL6J (Jackson Labs #000664) mice of both sexes (n=18). For inactivation experiments, parvalbumin (PV)-Cre knockin mice (Scholl et al. 2015) were crossed to a Cre dependent channelrhodopsin-2 (ChR2)-EYFP strain (Jackson Labs #024109, Madisen et al. 2012, n=10). These progenies selectively express ChR2 in PV+ interneurons. Mice were P35 and older to avoid the visual critical period. All animal procedures were approved by the University of Texas at Austin Institutional Animal Care and Use Committee.

#### Imaging

Experiments were conducted using 18 mice of both sexes older than P60. C57/BL6J was the primary background with up to 50% CBA/CaJ (Jackson Labs #000654)). GCaMP6 was expressed either transgenically (Ai162 [tm162.1(tetO-GCaMP6s,CAG-tTA2)Hze; Jackson Labs #031562] crossed to Slc17a7-IRES2-Cre-D [tm1.1(Cre)Hze; Jackson Labs #023527], n=10) or virally (Emx1-IRES-Cre [tm1(cre)Krj, Jackson Labs # 005628], n=3; Pvalb-IRES-Cre [tm1(cre)Arbr, Jackson Labs #017320], n=5). All animal procedures were approved by Duke University Institutional Animal Care and Use Committee.

### Physiology

#### Electrophysiology

Physiological procedures for mouse electrophysiology recordings are based on those previously described (Cang et al., 2008). Mice were anesthetized with 1,000 mg/kg urethane and 10 mg/kg chlorprothixene via intraperitoneal injection. A further intraperitoneal injection of 20 mg/kg dexamethasone was administered to prevent brain edema. During the course of the experiment, body temperature was monitored and maintained at 37oC. A tracheotomy was performed and the head was placed in a mouse adaptor (Stoelting). A small craniotomy (∼2-3 mm^2^) and durotomy were performed over the appropriate area of the sensory cortex. V1 was located by multiunit extracellular recordings with tungsten electrodes (1MΩ, Micro Probes). Mouse eyes were kept moist with a thin layer of silicone oil. The cortical surface was kept moist with saline or 4% agarose in normal saline.

We performed in-vivo whole cell recordings using the blind patch method. A silver-silver chloride wire was inserted into muscle near the base of the skull and used as a reference electrode. Pipettes (5-10 MΩ) were pulled from 1.2 mm outer diameter, 0.7 mm inner diameter KG-33 borosilicate glass capillaries (King Precision Glass) on a P-2000 micropipette puller (Sutter Instruments). Pipettes were filled with (in mM) 135 K-gluconate, 4 NaCl, 0.5 EGTA, 2 MgATP, 10 phosphocreatine disodium, and 10 HEPES, pH adjusted to 7.3 with KOH(Sigma-Aldrich). Neurons were recorded 150-500 μm below the cortical surface. Recordings were performed with a MultiClamp 700B patch clamp amplifier (Molecular Devices). Current flow out of the amplifier into the patch pipette was considered positive.

##### Cortical Inactivation

Inactivation experiments were restricted to layer IV, defined as 350-500 μm below the surface. A 470-nm fiber-coupled LED light was used to activate ChR2 in inhibitory parvalbumin positive neurons. The light covered the entire craniotomy and the light intensity was between 1 and 1.3 mW. The light was turned on 250 ms before the onset of visual stimulation.

#### Imaging

Animals were implanted with a titanium headpost and 5 mm cranial window as previously described (Goldey et al., 2014). Briefly, dexamethasone (3.2 mg/kg, s.c.) and Meloxicam (2.5 mg/kg, s.c.) were administered at least 2 h before surgery. Animals were anesthetized with ketamine (200 mg/kg, i.p.), xylazine (30 mg/kg, i.p.) and isoflurane (1.2-2% in 100% O2). Using aseptic technique, a headpost was secured using cyanoacrylate glue and C&B Metabond (Parkell), and a 5 mm craniotomy was made over the left hemisphere (center: 2.8 mm lateral, 0.5 mm anterior to lambda) allowing implantation of a glass window (an 8-mm coverslip bonded to two 5-mm coverslips (Warner no. 1) with refractive index-matched adhesive (Norland no. 71)) using Metabond.

The mice were allowed to recover for one week before habituation to head restraint. Habituation to head restraint increased in duration from 15 min to >2 h over 1-2 weeks. During habituation and imaging, mice were head restrained while allowed to freely run on a circular disc (InnoWheel, VWR).

Retinotopic maps were generated from GCaMP fluorescence or intrinsic autofluorescence. For intrinsic autofluorescence, the brain was illuminated with blue light (473 nm LED (Thorlabs) or a white light source (EXFO) with a 462 ± 15 nm band pass filter (Edmund Optics)), and emitted light was measured through a green and red filter (500 nm longpass). For GCaMP imaging, the same excitation light was used, but emitted light was measured through a 520 ± 18 nm band pass filter. For all conditions, images were collected using a CCD camera (Rolera EMC-2, Qimaging) at 2 Hz through a 5x air immersion objective (0.14 numerical aperture (NA), Mitutoyo), using Micromanager acquisition software (NIH). Images were analyzed in ImageJ (NIH) to measure changes in fluorescence (dF/F; with F being the average of all frames) to identify V1 and the HVAs, so as to target viral injections and/or imaging sessions to a monocular region of V1.

##### Viral injection

We targeted V1 in EMX1::Cre and PV::Cre mice for viral expression of GCaMP6s. Dexamethasone (3.2 mg/kg, s.c.) was administered at least 2 h before surgery and animals were anesthetized with isoflurane (1.2-2% in 100% O2). The coverslip was sterilized with 70% ethanol and the cranial window removed. A glass micropipette was filled with virus (AAV1.Syn.Flex.GCaMP6s.WPRE.SV40; titer: 1-3e13 GC/ml; UPenn CS1242), mounted on a Hamilton syringe, and lowered into the brain. 100 nL of virus was injected at 250 and 500 µm below the pia (30 nL/min); the pipette was left in the brain for an additional 10 minutes to allow the virus to infuse into the tissue. Following injection, a new coverslip was sealed in place. Imaging experiments were conducted at least one week following injection to allow for sufficient expression.

##### Two-photon imaging

Calcium imaging data was collected using a microscope controlled by Scanbox software (Neurolabware). Excitation light (920 nm) from a Mai Tai eHP DeepSee laser (Newport) was directed into a modulator (Conoptics) and raster scanned onto the brain with a resonant galvanometer (8 kHz, Cambridge Technology) through a 16X (0.8 NA, Nikon) water-immersion lens. Average power at the surface of the brain was 30-50 mW. Frames were collected either at 15 Hz (FOV of 680×605 µm) or at 30 Hz (415×735 µm). Emitted photons were directed through a green filter (510 ± 42 nm band filter; Semrock) onto GaAsP photomultipliers (H10770B-40, Hamamatsu). Images of excitatory cells were captured 193.7 ± 31.3 µm (range: 150-250 µm) below the pia, while inhibitory cells were imaged at 183.3 ± 12.4 µm (range: 150-225 µm).

##### Eye tracking

Pupil position was monitored via scattered infrared light from two-photon imaging. Light was collected using a GENIE Nano CMOS camera (Teledyne Dalsa) using a long-pass filter (695 nm) at the imaging frame rate.

### Stimulus Presentation

#### Electrophysiology

All stimuli were generated via the Psychophysics Toolbox (Brainard, 1997; Pelli, 1997) for MATLAB (Mathworks) on a Macintosh (Apple) computer. Stimuli were presented on a calibrated CRT monitor (Sony FDM-520) placed 25 cm in front of the animals’ eyes with a refresh rate of 100 Hz and a spatial resolution of 1,204 × 768 pixels. The mean luminance of the monitor was 40 cd/m^2^. Drifting gratings (at the preferred spatial frequency, temporal frequency of 2 Hz and variable diameter) were presented for a period of 4 seconds. Each stimulus was followed by a 1 second blank period. Test and mask gratings were presented at all combinations of four contrasts: 0%, 16%, 32% and 48%. For the phase shift experiments, the spatial phase of the test stimulus was fixed at 0 while the mask one of four phases: 0°, 90°, 180°, 270°. In a subset of neurons (n = 3) 8 spatial phases 45° apart were presented.

#### Imaging

Visual stimuli were presented on a 144-Hz (Asus) LCD monitor for imaging experiments. The monitor was calibrated with an i1 Display Pro (X-rite) for mean luminance at 50 cd/m^2^ and positioned 21 cm from the eye. All stimulus presentations used MWorks (https://mworks.github.io/) and custom software in MATLAB (MathWorks). All visual stimulus conditions within each experiment were randomly interleaved.

Visual stimuli were sinusoidal drifting gratings, 30 deg in diameter, placed in the monocular visual field. Four different visual stimulus protocols were used for the imaging experiments. A) “All directions plus mask”: Drifting gratings moving in 16 directions with and without an orthogonal mask (starting phase of 0º or 90º) at 50% contrast. Stimuli were presented for 1 s, with a 3 s mean luminance inter-trial interval (ITI), at a spatial frequency (SF) of 0.05 cycles/degree and a temporal frequency (TF) of 2 Hz. Data from 7 mice were collected from these experiments and are presented in Figures 1 and 7. B) “Two directions plus mask at 4 phases”: Drifting gratings moving rightward or upward each at three contrasts (0, 32 and 48%). The upward drifting grating was presented at 4 phases (0, 90, 180, or 270º). Stimuli were presented for 4 s, with a 4 s ITI, at a SF of 0.05 cycles/degree and a TF of 1 Hz. Data from 8 mice were collected from these experiments and are presented in Figure 5-6. C) “Two directions plus mask at 8 phases”: Drifting gratings moving rightward at 0 and either 32% or 48% contrast and upward at 0 and 48% contrast. The upward drifting grating was presented at 8 phases 45º apart. Stimulus duration, SF and TF were the same as in protocol B. These experiments were followed by a separate run in which we presented drifting gratings moving in 16 directions at matched SF and TF and 100% contrast. Data from 7 mice were collected from these experiments and are presented in Figures 5-7. D) “Stimulus adaptation”: Drifting gratings moving rightward or upward at each of five contrasts (0 6.25, 12.5, 25, 50%), either with or without adaptation. Adaptation was performed in blocks of 60 trials; adaptation blocks were preceded by a 2 min presentation of a 50% contrast, rightward drifting grating, followed by a 5 s top-up before each trial. The timing of the control block was the same, but the screen was at mean luminance. Test stimuli were presented for 1 s, at a SF of 0.1 cycles/degree and a TF of 2 Hz. Data from 7 mice were collected from these experiments and are presented in Figure 2.

### Feedforward Model

The model was adapted from a previous study (see details: Priebe & Ferster, 2006). Thalamic neurons were modeled as a single ON subregion for simplicity. To mimic the lower firing rates observed in mice, the firing rates were set to 7.3-11.3 Hz with a background firing rate of 0-2 Hz. The receptive fields of 8 thalamic relay cells were aligned along the preferred orientation of the neuron. To adjust the orientation selectivity of the model, the space between the receptive fields was systematically reduced (max = pi/4; min = 0). To simulate populations of model neurons, neurons were randomly assigned a value in this range. To mimic experimental conditions, the values of the test and mask phase were randomly assigned a value between 1° and 360°.

### Quantification and Statistical Analysis

#### Electrophysiology

Records were analyzed by using a 5-ms median filter to remove action potentials to isolate the membrane potential and spiking responses. Membrane potential and spiking responses were cycle averaged after removing the first cycle. We used the Fourier transform to calculate the mean and modulation amplitude for membrane potential and spiking responses.

#### Imaging

All two-photon imaging data was analyzed using custom code written in MATLAB (Mathworks). Image stacks from each imaging session were registered for x-y motion to the same stable reference image selected out of several 500-frame-average images, using Fourier domain subpixel 2D rigid body registration.

Cell bodies were manually segmented from the average change in fluorescence (dF/F) during stimulus presentation (where F is the average of 1 second preceding the stimulus) for each unique stimulus condition as well as the maximum projection across all stimulus conditions. Fluorescence time courses were generated by averaging all pixels in a cell mask. Neuropil signals were removed by first selecting a shell around each neuron (excluding neighboring neurons), estimating the neuropil scaling factor (by maximizing the skew of the resulting subtraction), and removing this component from each cell’s time course. Visually evoked responses were measured as the average dF/F during the stimulus presentation window, excluding the five frames immediately following stimulus onset to account for visual conduction delays. Cells were categorized as visually responsive if they had statistically significant responses to at least one stimulus condition compared to a similar baseline window, as measured with a one-sided t-test with the significance threshold Bonferroni corrected for the number of stimulus conditions. The preferred direction was determined by averaging across stimuli of the same direction and identifying the stimulus condition that drove the strongest response.

Masking index (MI) and selectivity index (SI) were only measured for conditions where the neuron responded significantly to either the test or the mask alone. For calculating MI, SI, and orientation selectivity index (OSI) responses below zero were set to zero, such that all values are constrained to be between 0 and 1. Cells with significant MI in the control condition (**Figure 2**) were determined with a two-way Student’s t-test comparing the distribution of plaid responses across trials to the sum of the average test and mask response.

##### Locomotion and eye tracking analysis

Locomotion was monitored with a digital encoder (US Digital) attached to a circular running wheel. In general, the mice were stationary and only ran (average speed greater than 2 cm/s) on a minority of trials (average: 3.6%, median: 0.8%; range 0-45%, 24 sessions). Thus, we combined all trials regardless of locomotion state.

We used pupil diameter as an alternative measure of arousal. Pupil size and position were extracted from each frame using the native MATLAB function imfindcircles, and quantified for each trial by averaging all frames during each stimulus presentation. To determine if there is a relationship between arousal and MI, we identified the trials from experiments from stimulus set C (since this had the smallest number of stimulus conditions) within the bottom (0.17 ± 0.01 mm, n = 7 mice) and top (0.27 ± 0.02 mm) quartile of pupil size. This increase in pupil diameter resulted in a significant increase in response to the gratings (bottom-0.19±0.01; top-0.34±0.02; n = 272 cells; paired t-test: p=9.9e-15), but no significant change in MI (bottom-0.03 ± 0.03; top--0.01 ± 0.03; p = 0.21).

To ensure that variable pupil position did not obscure the phase sensitivity of visual responses, average pupil position was quantified for these experiments (Figures 5-7). The average pupil position during each stimulus presentation was converted to degrees of visual angle with a 1:25 degrees to micrometer scale (Park et al., 2012). Only trials in which the pupil was within 2 deg of the median position were considered for analysis.

##### Phase sensitivity analysis

To measure the phase dependence, the masking index data were fit with a sinusoid:

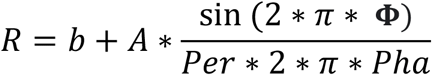

where b is the baseline offset, A is the amplitude, Per is the period, and Pha is the preferred phase. Shuffled fits were performed on data in which the phase-identity of trials was randomized. For electrophysiology data this was done on a cycle as opposed to a trial basis.

## Figure Legends

**Supplementary Figure 1.**
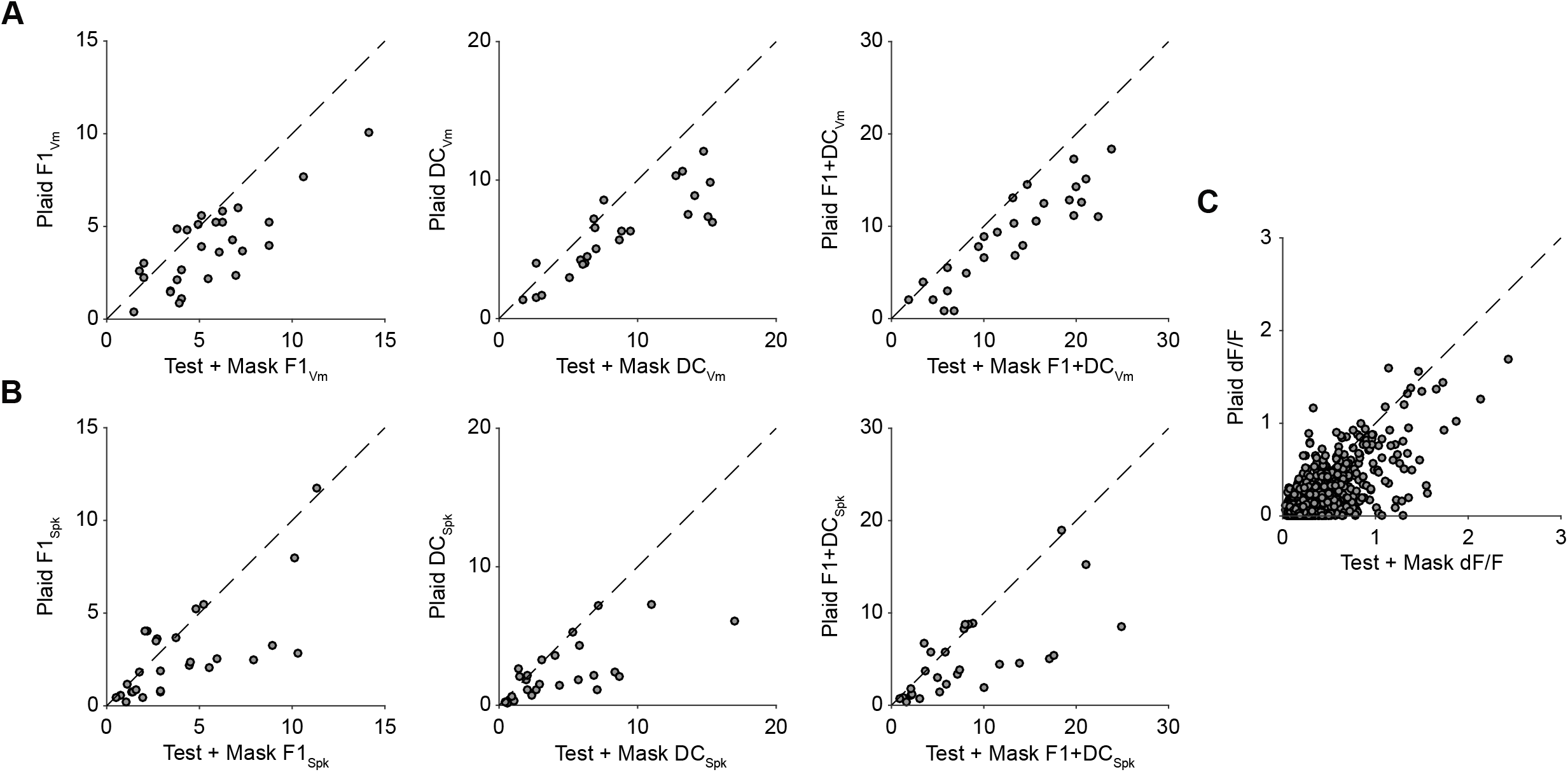
Effects of masking in mouse V1 **A**. Scatterplots comparing mean dF/F for test+mask and plaid. **B**. Scatterplots comparing membrane potential test+mask and plaid F1, DC and F1+DC. **C**. Same as (**B**) for spike rate.

**Supplementary Figure 2.**
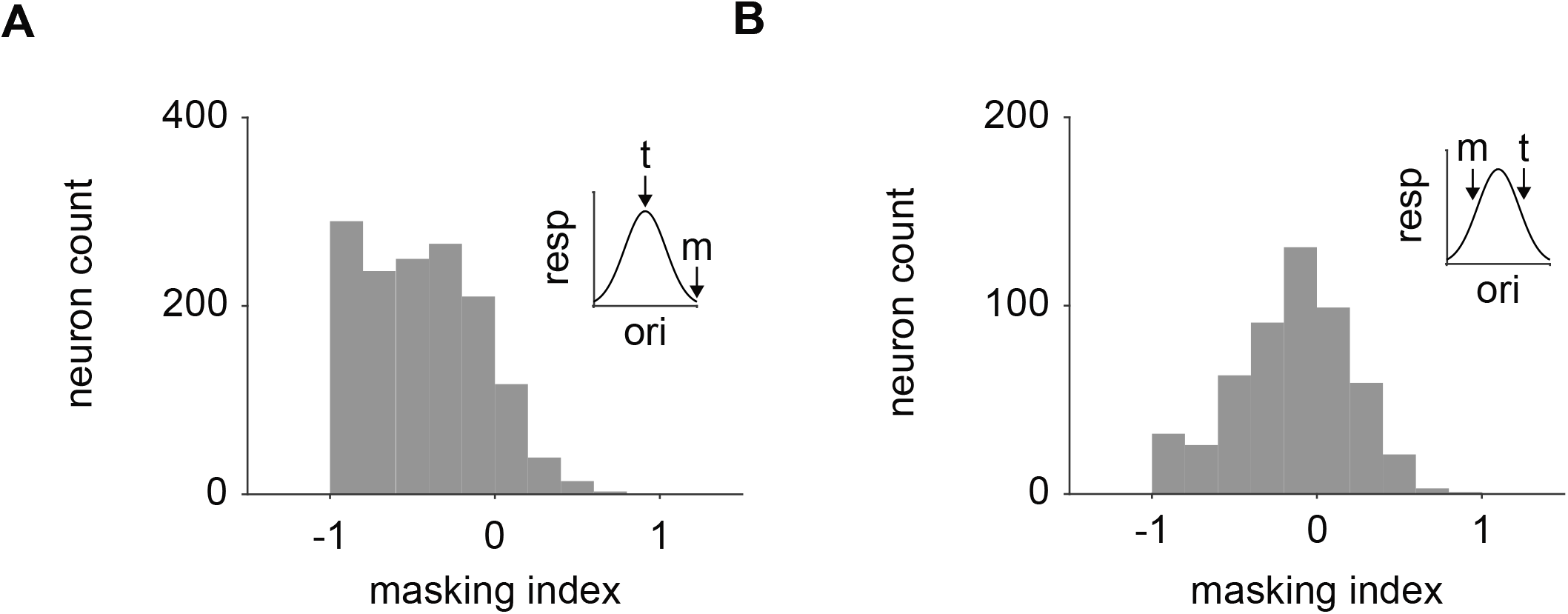
Effects of test stimulus orientation on masking **A**. Distribution of masking index from calcium imaging experiments when the test stimulus is the preferred orientation of the neuron. **B**. Same as **(A)** when the test is preferred orientation + 45°.

**Supplementary Figure 3.**
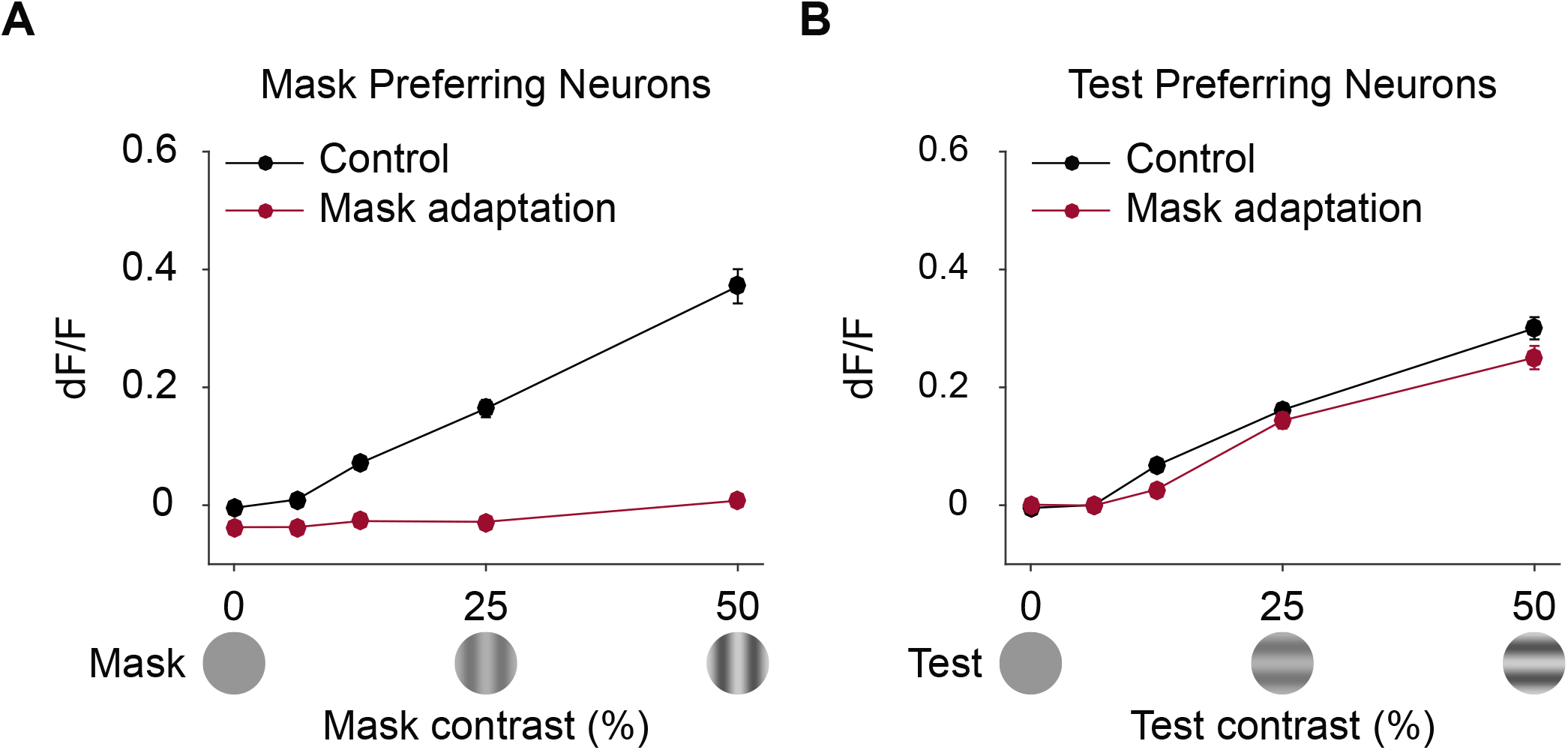
Effects of adaptation on responses to gratings **A**. Average dF/F in response to mask stimulus for neurons that preferred the mask stimulus. Responses are shown before (black trace) and after (red trace) adaptation to the mask. Error bars are standard error of the mean across cells. **B**. Same as **(A)** for responses to the test in neurons that prefer the test.

**Supplementary Figure 4.**
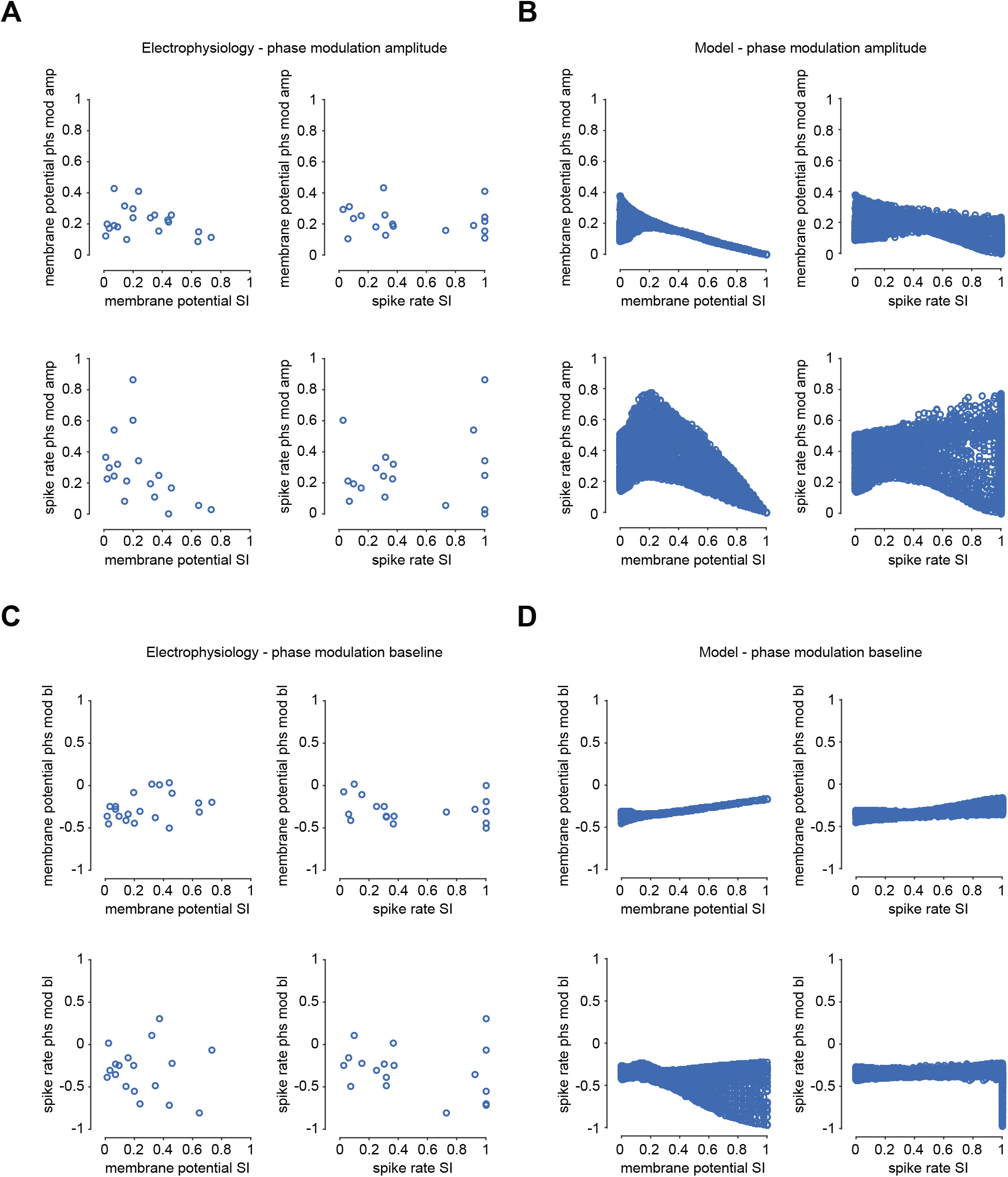
Relationship between selectivity index and phase modulation amplitude/baseline in electrophysiology data and model neurons **A**. Scatter plots comparing all combinations of the SI and phase modulation amplitude for spike rate and membrane potential data. **B**. Same as **(A)** but for model neurons. **C**. Same as **(A)** for phase modulation baselines. **D**. Same as **(C)** but for model neurons.

**Supplementary Figure 5.**
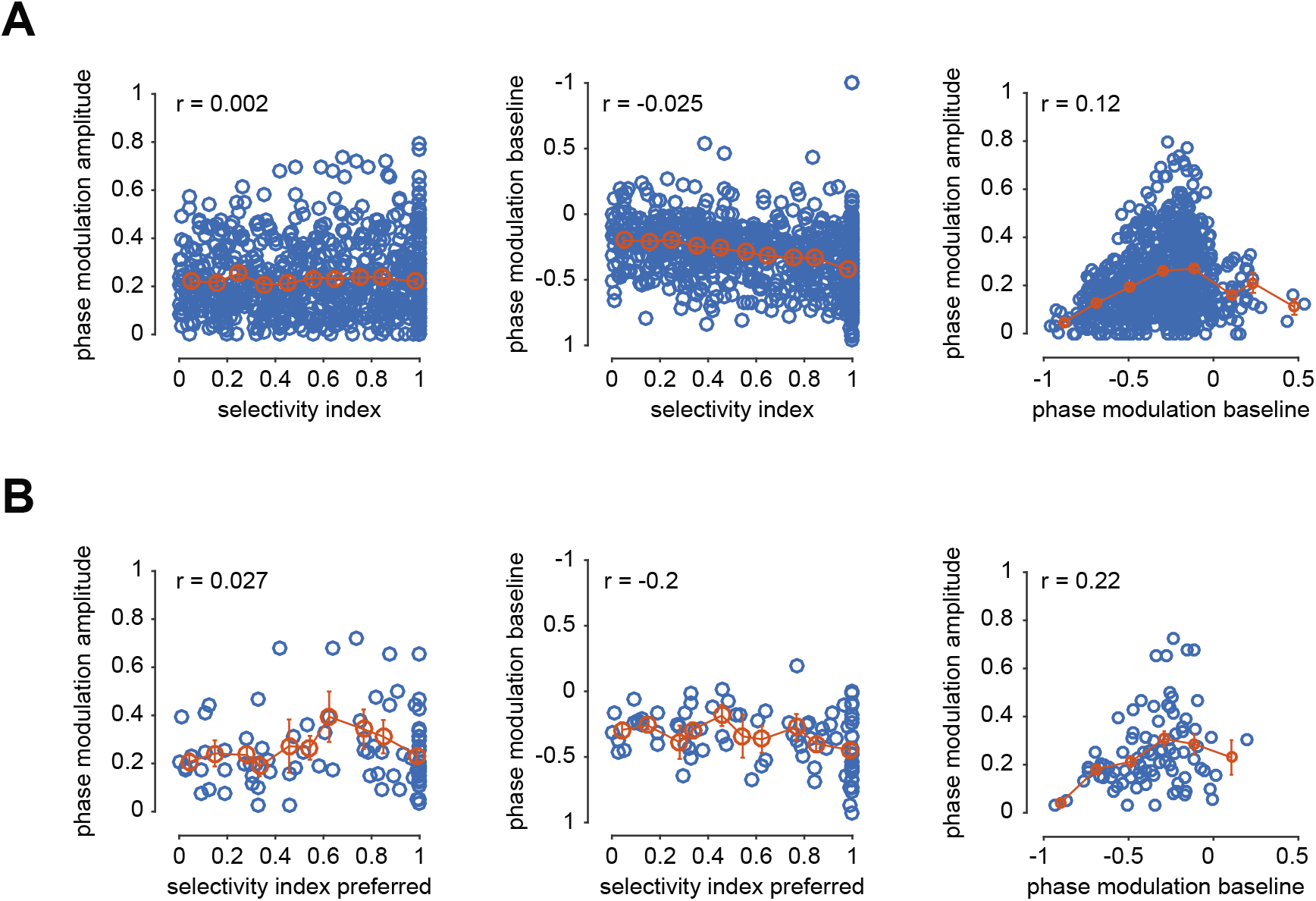
Relationship between selectivity index and phase modulation amplitude/baseline in calcium imaging data. **A**. Scatter plots comparing dF/F selectivity index (left), phase modulation amplitude (middle) and phase modulation baseline (right). Red circles are binned averages with error as standard error of the mean across cells. **B**. Same as **(A)** but for neurons where the selectivity index was measured at the preferred orientation.

